# Inhibition of the occipital face area modulates the electrophysiological signals of face familiarity positively: a combined cTBS-EEG study

**DOI:** 10.1101/2020.12.04.410035

**Authors:** Charlotta Marina Eick, Géza Gergely Ambrus, Gyula Kovács

## Abstract

The occipital face area (OFA) is hierarchically one of the first stages of the face processing network. It has originally been thought to be involved in early, structural processing steps, but currently more and more studies challenge this view and propose that it also takes part in higher face processing, such as identification and recognition. Here we tested whether the OFA is involved in the initial steps of recognition memory and plays a causal role in the differential processing of familiar and unfamiliar faces. We used an offline, inhibitory continuous theta-burst stimulation (cTBS) protocol over the right OFA and the vertex as control site. An electroencephalographic (EEG) recording of event-related potentials (ERPs), elicited by visually presented familiar (famous) and unfamiliar faces was performed before and after stimulation. We observed a difference in ERPs for famous and unfamiliar faces in a time-window corresponding to the N250 component. Importantly, this difference was significantly increased by cTBS of the right OFA, suggesting its causal role in the differential processing of familiar and unfamiliar faces. The enhancement occurred focally, at electrodes close to the right hemispheric cTBS site, as well as over similar occipito-temporal sites of the contralateral hemisphere. To the best of our knowledge, this is the first study showing the causal role of the rOFA in the differential processing of familiar and unfamiliar faces, using combined cTBS and EEG recording methods. These results are discussed with respect to the nature of familiar face representations, supported by an extensive, bilateral network.

## 1. Introduction

The recognition and identification of human faces are essential to our every-day life. While the exact details of these processes are still a subject of discussion, it seems that they can be divided into a few functionally and anatomically separate stages. First, familiarity refers to the vague feeling of “knowing” that an event or a person has already been encountered, without any specific associated knowledge. Second, recollection provides us additional episodic, contextual and semantic information about these events/persons and familiarity and recollection, together, are frequently referred to as recognition memory (Rugg & Yonelinas, 2003), a subtype of declarative memory. Finally, the determination of the facial identity of a person occurs.

The neurocognitive processes of the first step, underlying face familiarity representation, are best studied by comparing the neural responses to unfamiliar and familiar faces. It is widely accepted that there are significant perceptual as well as neural processing differences between the two (for reviews, see (Gobbini & Haxby, 2007; Johnston & Edmonds, 2009; Ramon & Gobbini, 2018; Young & Burton, 2017, 2018). For example, the matching of naturally variable images (i.e., “telling faces together”) is an easy and automatic task for familiar identities, while it is very difficult for unfamiliar ones (for a review, see Jenkins & Burton, 2011). This suggests that the specificity of the responsible neurons to low-level features, such as viewpoint or lightning depends strongly on face familiarity (for a review, see Johnston & Edmonds, 2009). Therefore, it is not surprising that numerous studies tested how the neural representation dynamics of familiar and unfamiliar faces differ, using electrophysiological (and magnetoencephalographical, MEG) methods (for a recent summary of the relevant neuroimaging studies see Kovács, 2020).

These studies found a difference in event-related potentials (ERPs) and MEG components for unfamiliar when compared to famous or personally familiar faces both in early (<170 ms) and late (>250 ms) time-windows (for reviews see Boehm et al., 2006; Gainotti, 2007; Huang et al., 2017; Ramon & Gobbini, 2018).

Studies testing the earliest occipito-temporal face-preferential ERP/MEG component (Bentin et al., 1996; Itier & Taylor, 2004), the N/M170 are inconsistent regarding its familiarity modulation. While some studies observed a decreased familiarity-related N/M170 (Jemel et al., 2003; Marzi & Viggiano, 2007; Todd et al., 2008) others reported enhancements for personally familiar or famous faces (Alonso-Prieto et al., 2015; Barragan-Jason et al., 2015; Caharel et al., 2002; Caharel et al., 2006; Caharel et al., 2014; Kloth et al., 2006; Wild-Wall et al., 2008), or no modulation at all (Keyes et al., 2010; Kotlewska & Nowicka, 2015).

Studies of the later components, on the other hand, reported more consistent effects. Regarding the N250, another occipitaltemporal component related to working memory representations of faces (for reviews see Schweinberger, 2011; Schweinberger & Neumann, 2016), most studies showed enhanced negativity for famous or personally familiar faces relative to unfamiliar ones (Andrews et al., 2017; Gosling & Eimer, 2011; Herzmann et al., 2004; Miyakoshi et al., 2008; Pierce et al., 2011; Tacikowski et al., 2011; Tanaka et al., 2006; Wiese et al., 2019; Wild-Wall et al., 2008; Wuttke & Schweinberger, 2019).

The last prominent ERP/MEG components which are associated to face processing are the centro-parietal N400 and P600, ranging from 300 - 800 ms post-stimulus onset and named collectively as late negativity (LN; Wuttke & Schweinberger, 2019). The LN shows lower (Bentin & Deouell, 2000; Caharel et al., 2005; Eimer, 2000; Herzmann & Sommer, 2010; Pfütze et al., 2002; Schweinberger et al., 2002; Wiese et al., 2019; Wuttke & Schweinberger, 2019) or higher (Henson et al., 2003) amplitudes for unfamiliar when compared to famous or personally familiar faces. In general, such differences of familiar and unfamiliar face ERP/MEG components (familiarity effect, FE) have been interpreted as the correlates of face-associated declarative memory and non-sensory, semantic knowledge activations (Huang et al., 2017).

But where do these familiarity-related electrophysiological differences originate from? Neurocognitive models traditionally connect face familiarity to earlier, modality-specific processing levels (Bruce & Young, 1986), corresponding to the “core” face processing regions of the occipital, fusiform and superior temporal areas (Gobbini & Haxby, 2007). In line with this theory, neuroimaging studies found that, indeed, all these areas show larger activity for famous or personally familiar when compared to unfamiliar faces (for reviews of famous faces see Natu & O’Toole, 2011 and of personally familiar faces see Ramon & Gobbini, 2018; Sugiura, 2014). Probing the role of the core network in the development of familiarity, in a series of transcranial magnetic stimulation (TMS) experiments, previously we demonstrated that the right OFA (rOFA) plays a causal role in the experimental formation of familiarity (Ambrus, Windel et al., 2017) as well as in familiarity decisions for famous faces in visual (Ambrus, Dotzer et al., 2017) and cross-domain priming paradigms (Ambrus et al., 2019). These studies suggest that the role of rOFA extends over the traditionally proposed low-level, early, “entry-point” towards higher-level, image-invariant identity processing.

In the present study, we aimed at testing explicitly how the inactivation of the rOFA by continuous theta-burst TMS (cTBS; Huang et al., 2005) modulates the electrophysiological correlates of face familiarity. cTBS is an inhibitory TMS protocol that has previously been shown to inhibit visual cortical and motor cortex excitability as well as cognitive functions (Boroojerdi et al., 2000; Cao et al., 2018; Chen et al., 1997; Evers et al., 2001; Jing et al., 2001; Muellbacher et al., 2000; Romero et al., 2002; Rossi et al., 2000; Touge et al., 2001). We applied an offline cTBS design, where the electrophysiological signals are analyzed before as well as after the temporally separate application of TMS pulses. Such designs have previously been shown to modulate ERPs (Bohotin et al., 2002; Cao et al., 2018; Enomoto et al., 2001; Evers et al., 2001; Fumal et al., 2003; Jing et al., 2001; Rossi et al., 2000; Sadeh et al., 2011; Thut et al., 2003) and therefore they are ideal for establishing the causal role of an area in the emergence of a given ERP-related phenomenon (for a review see Pitcher et al., 2020). As previous studies revealed inhomogeneous results regarding the effect of cTBS on cortical excitability, the interpretation of reduced or enhanced ERP absolute amplitude values is difficult (for a review see Jost et al., 2020). Therefore here, instead of comparing pre and post cTBS-ERPs, we concentrated on the differential processing of familiar and unfamiliar faces, expressed in the [familiar-unfamilar] difference of ERP component amplitudes, after cTBS of the rOFA or Vertex, as control area. We hypothesized that if the rOFA plays a role in the sensation of face familiarity then the electrophysiologically observed FE will be modulated by the cTBS of the area. We reasoned that if cTBS reduces the differences of [familiar-unfamilar] ERPs then one can assume that the rOFA plays a positive role in the emergence of familiarity representation. If, however, the [familiar-unfamilar] ERP differences are enhanced by cTBS, it likely signals the onset of secondary, top-down mechanisms, compensating for inhibition (Thut & Pascual-Leone, 2010). In line with previous studies of face familiarity, we observed that cTBS of the rOFA affected the ERP familiarity effect around the time-window of the N250 component the most. cTBS enhanced [familiar-unfamilar] ERP differences, thereby supporting further the important role of rOFA in face familiarity processing and suggesting the existence of compensatory, top-down mechanisms during its inhibition.

## 2. Methods

### 2.1 Participants

In total, 24 right-handed participants with normal or corrected to normal vision were tested (23.48 ± 2.91 (mean age ± SD), 2 male). The sample size was based on similar previous studies evaluating the OFAs role in face processing (Amado et al., 2018; Ambrus, Dotzer et al., 2017; Ambrus, Windel et al., 2017; Eick et al., 2020). Participants reported no prior neurological disorders or medication intake relevant to this study. Informed written consent was given before participation and the experimental procedures were approved by the Ethics Committee of Friedrich-Schiller University Jena, conforming to the Declaration of Helsinki. Participants received partial course credits or monetary compensation.

### 2.2 Localization of OFA

The right OFA was localized individually, via functional neuroimaging measurements prior to the first EEG session. A 3T magnetic resonance imaging (MRI) scanner (Siemens MAGNETOM Prisma fit, Erlangen, Germany), at the Institute for Diagnostic and Interventional Radiology, University of Jena was used for the experiments. Technical specifications for the structural T1-weighted images were as follows: MPRAGE sequence; TR = 2300 ms; TE = 3.03 ms; isotropic voxel size = 1 mm³. For the functional MRI images, a 20-channel phased-array head coil and a gradient-echo, T2-weighted EPI sequence were used (35 slices, 10° tilted relative to axial, TR = 2000 ms, TE = 30 ms, flip angle = 90°, matrix array = 64 × 64 and isotropic voxel size = 3 mm³). The paradigm, applied during the functional scanning was implemented in Matlab (MathWorks, Version R2013). It included five blocks, each showing three, 20-second-long blocks of colored images of famous and unfamiliar faces, objects and Fourier-randomized noise patterns (Dakin et al. 2002), interrupted by 20 s long resting periods. The stimulus presentation frequency was 4 Hz (230 ms exposition time with a 20 ms inter-stimulus interval). The analysis of the data, including pre-processing and univariate statistics, was performed by SPM12 (Wellcome Department of Imaging Neuroscience, London, UK, Version 12). To determine the location of the rOFA the local maximum from the t-maps with a threshold of p < .05 _FWE_ or p < .0001 _uncorrected_ were chosen, by contrasting faces vs. objects and noise patterns. The average (± SE) MNI coordinates were 42.8 (3.48), − 79 (5.77), - 11 (4.77) for the right OFA (Fig. 1). Due to technical failures during scanning, the stimulation coordinates for one female participant were based on the mean of 60 coordinates of the rOFA, collected from female participants during previous studies of our laboratory.

**Fig. 1.**
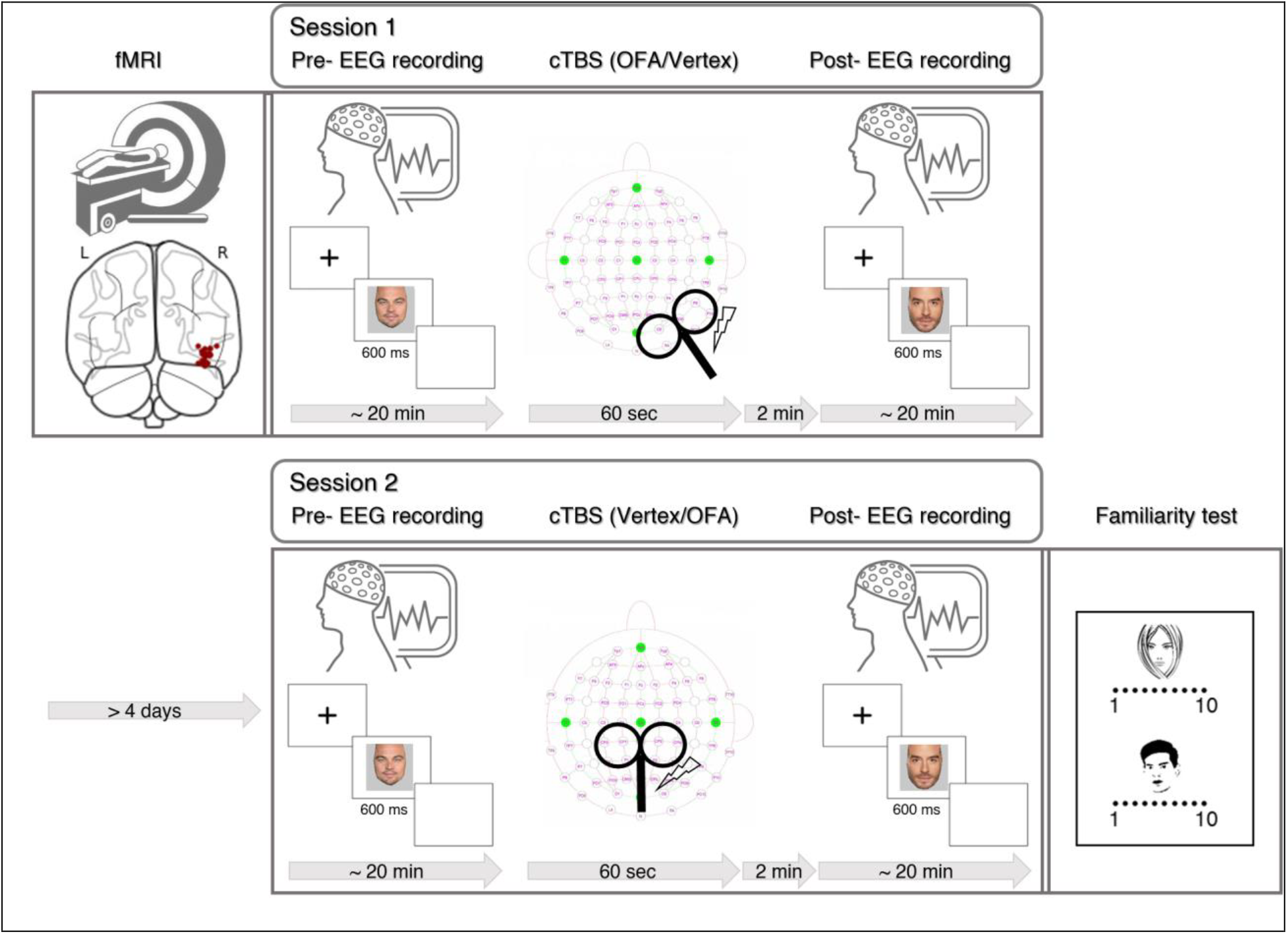
Experimental paradigm. The study began with an fMRI measurement to determine the location of the right OFA. The coronal section shows the OFA locations schematically. Session 1 included an EEG recording measurement, followed by a cTBS stimulation over either the rOFA or Vertex (randomized across participants) and a post-stimulation EEG recording. After a minimum of four days, a second session took place with another pre-TMS EEG measurement, cTBS stimulation of the other cortical region, another post-TMS EEG recording. Finally, a familiarity rating test was performed by the participants.

### 2.3 Stimuli

Faces of ten famous male individuals were used as stimuli. Five of these were celebrities, very familiar to our native German participants (Leonardo DiCaprio, Brad Pitt, Elyas M′Barek, Till Schweiger, Klaas Heufer-Umlauf), while another five were unknown and unfamiliar to them (Mika, Philippe Bas, Arjen Lubach, Garou, Juan Pablo Espinosa). Twenty-two colored photographs per identity were selected from the world wide web, showing frontal faces with positive or neutral expressions. The images were adjusted for brightness and cropped so that only the inner facial features were visible. The eye-levels were aligned and the distance between the eyes was matched to 118 - 127 pixels (see Ehrenberg et al., 2020). Two sets of 10 images per identity plus two target images for the target trials (see below) were used.

### 2.4 Experimental Procedure and EEG acquisition

The 110 images (10 faces + one target face of 10 individuals) were presented in a randomized order during EEG recordings. Each image was repeated 6 times, leading to 660 trials, separated into five runs (22 min recording time). Stimulus exposition time was 600 ms with an interstimulus interval of 1550 ms.

Face images were presented on a grey background with a central fixation cross on a TFT display (1680 × 1050 pixel resolution; refresh rate 60 Hz), using PsychoPy (Peirce, 2007). Participants were instructed to react with a button press when the target image appeared, which was tilted by 10 degrees either towards the right or left (10 % of trials).

After the completion of the second experimental session (Fig. 1), a detailed questionnaire was filled out by the participants, in which familiarity of each ID was evaluated on a Likert-scale from 1 (unfamiliar) to 10 (familiar). The results of this questionnaire suggest that the five familiar faces were indeed significantly more familiar to our participants than the five unfamiliar faces (ratings for familiar faces: 9.6 ± 0.7; ratings for unfamiliar faces: 2.8 ± 1.3; paired t-test for familiarity ratings: t(23) = 24.64, p > .001).

EEG was recorded by using a 64-channel Biosemi Active II system and ActiView (Version 6.05). Electrode impedances were kept below 10 kΩ and the bandwidth was 50 Hz (direct current DC) to 120 Hz. The sampling rate was 512 Hz and channels CMS and DRL served as ground and reference, respectively in a Biosemi introduced feedback loop, which reduces impedance. Recordings took place in an electrically and sound shielded and dimmed chamber. The eye-stimulus distance was 96 cm and reduced head and face movement were achieved by a chin rest. The experiment consisted of two sessions, separated by at least four days, one for stimulating the rOFA and another for the vertex, as control area. The order of these sessions was randomized and counterbalanced. During each session, first the EEG was recorded, then the cTBS was performed, immediately followed by the second EEG recording. The two EEG measurements contained different sets of images. The electrodes over the stimulated areas were removed from the EEG cap, prior to cTBS and replaced immediately afterward.

### 2.5 Transcranial magnetic stimulation

TMS was carried out using a PowerMag 100 Research Stimulator (MES Forschungssysteme GmbH). The neuronavigation system PowerMag View (MES Medizintechnik GmbH) was applied to guide the center of the stimulation cone to the rOFA or Vertex. The stimulation coil was handheld by the experimenter with the handle pointed backward. A chin rest stabilized the participants’ head during stimulation.

A continuous theta-burst stimulation (cTBS, 300 trains á 3 pulses (50 Hz; 900 total) with a 200 ms inter-train interval (5 Hz)) was delivered for 60 seconds, continuously at a maximal stimulation intensity of 40 % (Chiou & Lambon Ralph, 2018). This protocol has previously been shown to elicit a long-lasting (approximately 20 min post-stimulation time; Brückner et al., 2013; Huang et al., 2005; Tarnutzer et al., 2013) inhibitory effect over the underlying stimulated areas. Therefore, the second, post-cTBS EEG measurements remained within the time period of the expected cTBS aftereffects (average time between the end of the stimulation and the beginning of the EEG recording was 2.1 ± 0.1 minutes with an average duration of the post-EEG recording of 23.1 ± 0.1 min.).

### 2.6 Electrophysiological analysis

A Butterworth Bandpass Filter between 0.1 Hz and 40 Hz was applied and all channels were re-referenced by the averaged activity of all electrodes. The epochs were segmented from −200 ms to 850 ms, relative to stimulus onset and baseline correction was implemented for the timeframe −200 ms to 0 ms. Artifact rejection was performed by autocorrecting or when no interpolation was possible, auto rejecting bad channels, using the algorithm of Jas et al. (2017). The excluded data did not exceed the amount of 25 % of the trials (except for one participant, whose data was therefore excluded from further statistical analysis). The data were downsampled with a ratio of 2 to 256 Hz to decrease data size and to increase the signal-to-noise ratio (Grootswagers et al., 2017).

ERPs were then averaged for familiar and unfamiliar faces and each electrode and participant separately. The N170 (130-180 ms), N250 (200-400 ms) and LN (400-800 ms) ERP components were measured, according to the component peaks, identified on the grand-averages across conditions. The data analysis was performed in Python via Spyder, using MNE, pandas and NumPy packages (see Gramfort et al., 2013; Gramfort et al., 2014).

The familiarity effect was quantified on the electrodes, showing maximal ERP component amplitudes and corresponding to those identified in previous studies (Bindemann et al., 2008; Schweinberger et al., 2002; Schweinberger et al., 2004; Wiese et al., 2019; Wuttke & Schweinberger, 2019). Therefore, the electrodes of interest included PO8, PO10 and P8, P10 which lay in the proximity of the stimulating position, as well as the corresponding left hemisphere electrodes (PO7, PO9, P7, P9).

For the statistical analysis, we first tested if the N250 and LN ERP components reflect the differences of face familiarity, as suggested by previous research (Schweinberger, 2011; Schweinberger & Neumann, 2016; Wiese et al., 2019; Wuttke & Schweinberger, 2019). For this analysis, we used the data from the first EEG measurement, recorded prior to cTBS. As recent evidence suggests that averaging ERP amplitudes across electrodes increases the quality of the signal (Huffmeijer et al., 2014) the peak amplitudes were averaged across the electrodes of interest of the right and left hemispheres, separately for familiar and unfamiliar faces. Next, we compared the ERP amplitudes for familiar and unfamiliar faces of these two clusters, by a 2 × 2 repeated-measures ANOVA with the face familiarity (unfamiliar and familiar) and hemisphere (left and right) as factors.

Second, for evaluating the effect of cTBS on familiarity processing we ran a 2 × 2 repeated-measures ANOVA with cTBS location (rOFA and Vertex) and face familiarity (unfamiliar and familiar) as factors. Here we aimed for a higher spatial resolution with ERPs, so we performed this analysis step for each of the eight electrodes of interest separately.

## 3. Results

### 3.1 Pre-TMS familiarity effect is manifest on the N250

Familiar and unfamiliar faces elicited significantly different ERPs in the pre-cTBS recordings (Fig 2 a, b). The scalp topographies of the differences in ERP between familiar and unfamiliar faces were in line with previous results and suggested that this familiarity effect is concentrated over bilateral occipito-temporal areas. The effect started at around 200 ms post-stimulus onset and lasted up to 400 ms, corresponding to that of Wiese et al. (2019) for personally familiar faces (Fig 2 c). The ANOVA on the ERP amplitudes (Fig 2 d) revealed that familiarity affected N250 similarly for the two hemispheres (main effect of familiarity: F (1,22) = 7.14, p = .014, η^2^_p_ = .245; interaction of hemisphere x familiarity: F (1,22) = .83, p = .37, η^2^_p_ = .036).

**Fig 2.**
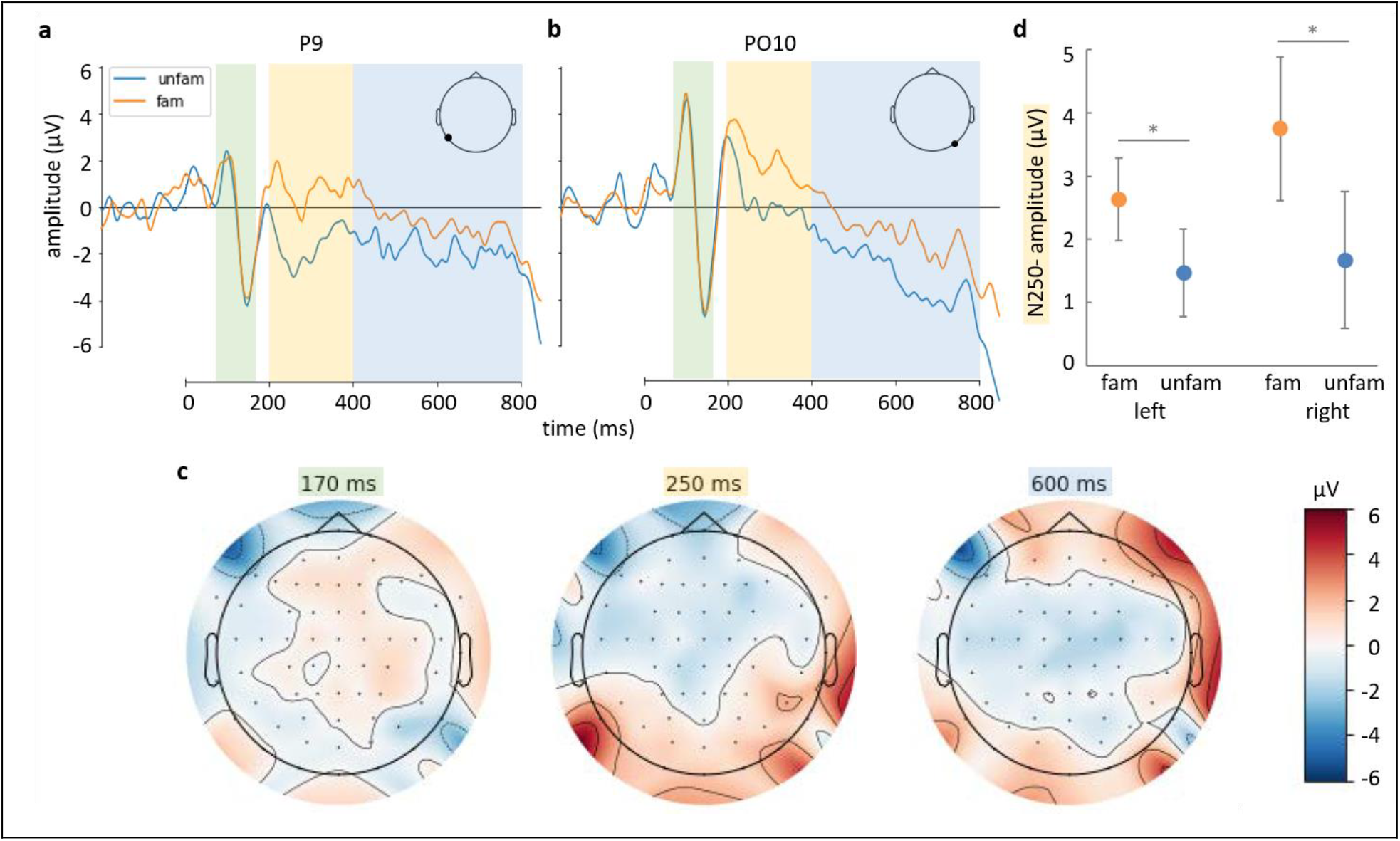
Familiarity effects on the ERPs. Grand average ERPs of a left **(a)** and right **(b)** electrodes, separately for familiar (orange) and unfamiliar (blue) faces. Shadings indicate the time windows of N170 (green), N250 (yellow), as well as the LN (blue) time windows. **(c)** Scalp-topographies for the [familiar-unfamiliar] difference at three representative time-windows (average of ± 15 ms). **(d)** The averaged amplitudes of the N250 component over the left and right electrode clusters for familiar (orange) and unfamiliar (blue) faces (mean ± SE; * ≤ .05).

### 3.2 cTBS of the rOFA enhances familiarity effect

Table I summarizes the main results of the statistical analysis for the N170, N250 and LN time-windows. As the table shows, similarly to the pre-TMS EEG measurements, familiarity led to different ERPs in a time-window, corresponding to the N250 (Fig. 3 a, c). This effect was significant over the PO8, PO10, P8, as well as over the PO7 and P7 electrodes, but all other electrodes showed a strong tendency for FE as well. Importantly, this FE was superseded by a significant interaction of TMS stimulation in one channel (PO8; Fig. 3 c). To our surprise, the cTBS x familiarity interaction was also significant over the non-stimulated, left hemisphere for one channel (P9; Fig. 3 a). While the analysis of the LN component revealed no significant effects, it is worth mentioning that the same electrode of the right hemisphere (PO8) showed a strong tendency for familiarity effect, as well as for its interaction with cTBS in the time-window of LN, too. Overall, cTBS interacted with familiarity-dependent processing in a few channels, rather than being distributed across the occipito-temporal cortex and the scalp topographies also show this spatially restricted interaction (Fig. 3 b). To test further, if the cTBS of the right hemisphere OFA indeed changes the N250 bilaterally we also estimated the scalp topographies of the [OFA - Vertex] difference, separately for unfamiliar and familiar faces (Fig. 3 e). These topographies show clearly, that indeed, cTBS over the right OFA altered the N250 ERP component over both hemispheres, specifically for the familiar faces, suggesting the close functional connection of the two hemispheres during their processing.

**Table I.** Summary of the statistical analysis after cTBS for each electrode of interest and time-window, separately. Familiarity: Main effect of familiarity (fam- unfam); cTBS location: Main effect of cTBS location (rOFA- Vertex); Interaction: Interaction of cTBS location and familiarity effects. Dark grey shadings indicate electrodes and conditions with significant results, while light grey shadings indicate strong tendencies.

3.2 cTBS of the rOFA enhances familiarity effect
|  |  | Familiarity | cTBS location | Interaction |
| --- | --- | --- | --- | --- |
| **N170** |  |  |  |  |
| Left | P7 | $F(1,22) = 2.49, p = .129, \eta^2_p = .102$ | $F = 0.45, p = .511, \eta^2_p = .020$ | $F = 0.02, p = .901, \eta^2_p = .001$ |
| | P9 | $F(1,22) = 3.27, p = .084, \eta^2_p = .129$ | $F = 0.39, p = .537, \eta^2_p = .018$ | $F = 0.36, p = .557, \eta^2_p = .016$ |
| | PO7 | $F(1,22) = 0.06, p = .812, \eta^2_p = .003$ | $F = 1.36, p = .256, \eta^2_p = .058$ | $F = 0.00, p = .971, \eta^2_p = .000$ |
| | PO9 | $F(1,22) = 0.60, p = .448, \eta^2_p = .026$ | $F = 1.25, p = .276, \eta^2_p = .054$ | $F = 0.46, p = .507, \eta^2_p = .020$ |
| Right | P8 | $F(1,22) = 3.26, p = .085, \eta^2_p = .129$ | $F = .528, p = .475, \eta^2_p = .023$ | $F = 1.61, p = .218, \eta^2_p = .068$ |
| | P10 | $F(1,22) = 0.26, p = .612, \eta^2_p = .012$ | $F = 0.21, p = .651, \eta^2_p = .009$ | $F = 1.88, p = .185, \eta^2_p = .079$ |
| | PO8 | $F(1,22) = 1.37, p = .255, \eta^2_p = .059$ | $F = 0.01, p = .923, \eta^2_p = .000$ | $F = 3.38, p = .079, \eta^2_p = .133$ |
| | PO10 | $F(1,22) = 5.30, p = .031, \eta^2_p = .194$ | $F = 1.06, p = .314, \eta^2_p = .046$ | $F = 1.56, p = .225, \eta^2_p = .066$ |
| **N250** |  |  |  |  |
| Left | P7 | $F(1,22) = 5.68, p = .026, \eta^2_p = .205$ | $F = 0.00, p = .952, \eta^2_p = .000$ | $F = 1.31, p = .264, \eta^2_p = .056$ |
| | P9 | $F(1,22) = 3.83, p = .063, \eta^2_p = .148$ | $F = 1.17, p = .291, \eta^2_p = .051$ | $F = 4.68, p = .042, \eta^2_p = .175$ |
| | PO7 | F (1,22) = 4.89, p = .038, $\eta^2_p$ = .182 | F = 0.18, p = .672, $\eta^2_p$ = .008 | F = 0.80, p = .382, $\eta^2_p$ = .035 |
| | PO9 | F (1,22) = 2.34, p = .140, $\eta^2_p$ = .096 | F = 0.06, p = .806, $\eta^2_p$ = .003 | F = 2.47, p = .130, $\eta^2_p$ = .101 |
| Right | P8 | F (1,22) = 8.54, p = .008, $\eta^2_p$ = .280 | F = 1.16, p = .293, $\eta^2_p$ = .050 | F = 1.59, p = .220, $\eta^2_p$ = .067 |
| | P10 | F (1,22) = 2.61, p = .120, $\eta^2_p$ = .106 | F = 1.00, p = .329, $\eta^2_p$ = .043 | F = 2.21, p = .152, $\eta^2_p$ = .091 |
| | PO8 | F (1,22) = 4.35, p = .049, $\eta^2_p$ = .165 | F = 0.05, p = .822, $\eta^2_p$ = .002 | F = 4.19, p = .05, $\eta^2_p$ = .160 |
| | PO10 | F (1,22) = 5.07, p = .035, $\eta^2_p$ = .187 | F = 3.42, p = .078, $\eta^2_p$ = .134 | F = 1.13, p = .300, $\eta^2_p$ = .049 |
| **LN** |  |  |  |  |
| Left | P7 | F (1,22) = 2.67, p = .116, $\eta^2_p$ = .108 | F = .506, p = .484, $\eta^2_p$ = .022 | F = 1.21, p = .284, $\eta^2_p$ = .052 |
| | P9 | F (1,22) = 1.20, p = .286, $\eta^2_p$ = .052 | F = 0.00, p = .970, $\eta^2_p$ = .000 | F = 1.35, p = .258, $\eta^2_p$ = .058 |
| | PO7 | F (1,22) = 1.19, p = .288, $\eta^2_p$ = .051 | F = 0.07, p = .794, $\eta^2_p$ = .003 | F = 0.96, p = .337, $\eta^2_p$ = .042 |
| | PO9 | F(1,22) = 1.18, p = .289, $\eta^2_p$ = .051 | F = 0.05, p = .831, $\eta^2_p$ = .002 | F = 0.85, p = .366, $\eta^2_p$ = .037 |
| Right | P8 | F (1,22) = 1.05, p = .316, $\eta^2_p$ = .046 | F = 4.15, p = .054, $\eta^2_p$ = .159 | F = 2.88, p = .104, $\eta^2_p$ = .116 |
| | P10 | F (1,22) = 0.42, p = .524, $\eta^2_p$ = .019 | F = 2.87, p = .104, $\eta^2_p$ = .115 | F = 1.19, p = .286, $\eta^2_p$ = .051 |
| | PO8 | F (1,22) = 3.45, p = .077, $\eta^2_p$ = .135 | F = 0.16, p = .692, $\eta^2_p$ = .007 | F = 3.23, p = .086, $\eta^2_p$ = .128 |
| | PO10 | F (1,22) = 2.23, p = .149, $\eta^2_p$ = .092 | F = 1.02, p = .323, $\eta^2_p$ = .044 | F = 0.26, p = .616, $\eta^2_p$ = .012 |

**Fig. 3.**
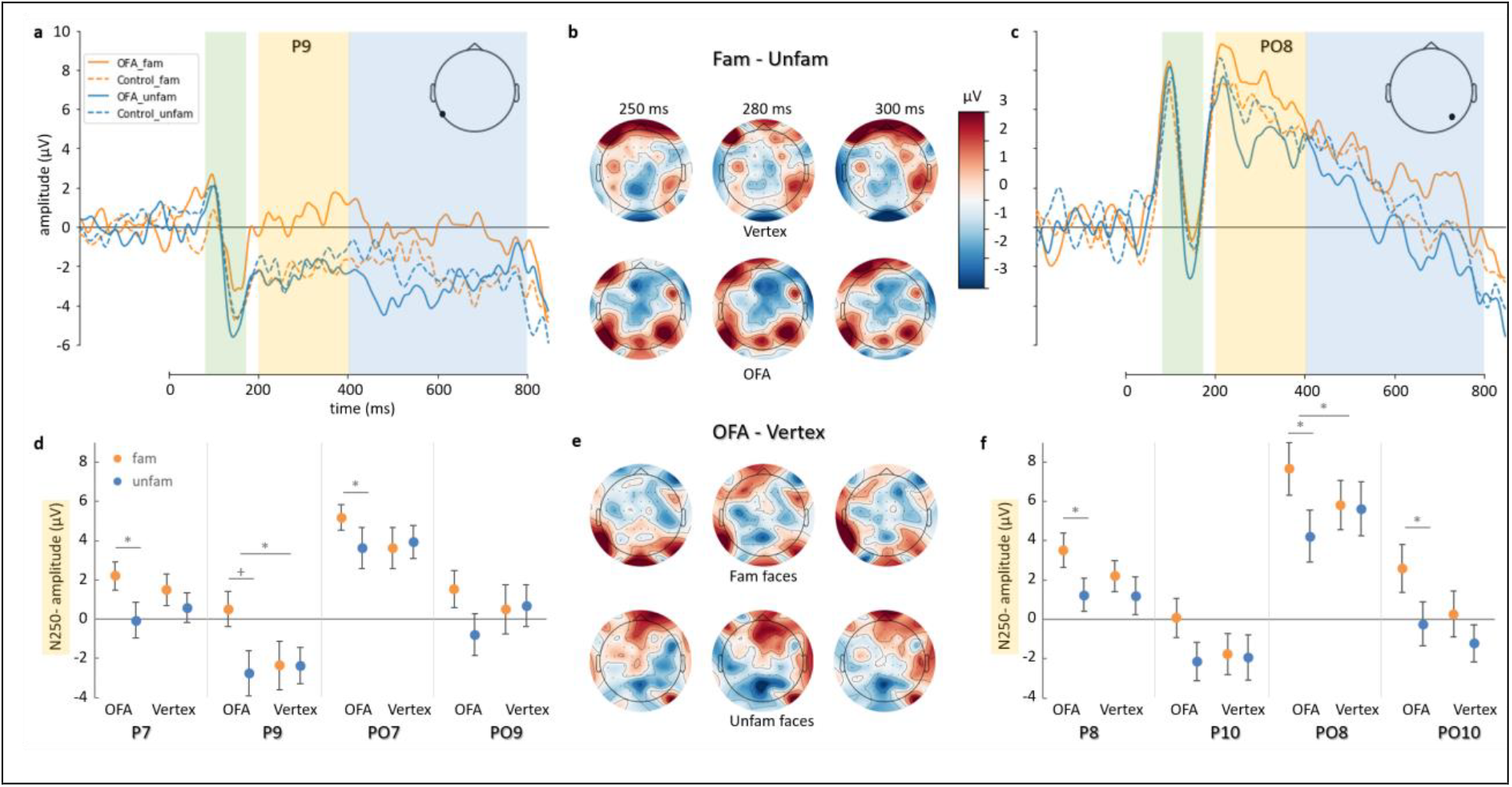
Grand average ERPs of P9 **(a)** and PO8 **(c)** channels, separately for familiar (orange) and unfamiliar (blue) faces, after cTBS to the rOFA (continuous lines) and the vertex (dotted lines). **(b)** Scalp-topographies for the [familiar-unfamiliar] difference for the rOFA and Vertex stimulation, for 250, 280 and 300 ms post stimuli onsets (average of ± 15 ms). The peak amplitudes of the N250 component over the left **(d)** and right electrode channels **(f)** for familiar (orange) and unfamiliar (blue) faces (mean ± SE; * ≤ .05, + ≤ .063). **(e)** Scalp-topographies for the [OFA - Vertex] difference for familiar and unfamiliar faces, for 250, 280 and 300 ms post stimuli onsets (average of ± 15 ms).

## 4. Discussion

To the best of our knowledge, this is the first combined TMS-EEG study, testing the causal role of the rOFA in the differential processing of un/familiar faces. The major result of our study is that the inactivation of the right OFA by continuous theta-burst TMS increases the difference between familiar and unfamiliar face evoked ERPs in a time-window, corresponding to the N250 component. The enhancement is not distributed over the occipito-temporal cortex, rather it occurs focally, at electrodes close to the right hemispheric TMS site, as well as over similar occipito-temporal sites of the contralateral hemisphere.

### 4.1 The N250 component shows familiarity effects

Our current results confirm those prior studies which showed that the earliest face familiarity sensitive ERP component is the N250, both for pre-experimental as well as for newly acquired familiarization states (Andrews et al., 2017; Gosling & Eimer, 2011; Huang et al., 2017; Schweinberger et al., 2002; Schweinberger et al., 2004; Tanaka et al., 2006, but see Alonso-Prieto et al., 2015 for earlier correlates). We used naturally variable faces of celebrities both as familiar and unfamiliar faces, comparable to the stimuli of the recent study of Wiese et al. (2019). Interestingly, the results of that paper differ from ours in two crucial aspects. First, while our study, similarly to several prior ones, showed familiarity effects for famous faces on the N250 (Gosling & Eimer, 2011; Huang et al., 2017), as well as in later time-periods, corresponding to LN (Bentin & Deouell, 2000; Eimer, 2000; Henson et al., 2003), no differences of N250 or LN amplitudes were reported by Wiese et al. (2019) for famous and unfamiliar faces. Second, while those studies which previously found FE on the N250 and subsequent ERP components typically reported more negative ERP amplitudes for familiar relative to unfamiliar faces, in our study we found the effect in the opposite direction. Our results are in correspondence with the findings of Henson et al. (2003), however, who found more positive amplitudes for familiar when compared to unfamiliar faces in a late time-window (from 550 ms onwards). We argue that the direction of the [familiar-unfamiliar] ERP difference is not crucial to the aims of the current study for several reasons. First, the interpretation of the ERP wave directions is complicated by the fact that both, the depolarization of apical dendrites in superficial cortical layers, as well as the hyperpolarization of deep cortical layer neurons manifest themselves on the scalp as a negative ERP deflection (Kotchoubey, 2006). Second, in our current experiment, we concentrated on the modulatory effect of cTBS on the [familiar-unfamiliar] ERP differences. Nonetheless, either future systematic studies, manipulating the level of familiarity, together with image properties and tasks, or preferably multivariate pattern analysis (MVPA) techniques, decoding the available information about familiarity are necessary to confirm the exact temporal dynamics and directions of these effects. Indeed, a few recent studies applied MVPA to EEG/MEG data and found that face familiarity relies on a cascade of neural processes, ranging from 85 ms post-stimulus onset and lasting up until 600 ms (Barragan-Jason et al., 2015; Dobs et al., 2019; Ehrenberg et al., 2020), paving a very promising, novel path towards understanding person recognition and identification.

### 4.2 The role of OFA in higher-level face processing stages

The N250, as well as the repetition sensitive N250r (Begleiter et al., 1995; Schweinberger et al., 1995), are currently considered as indicators of early, probably perceptual stages of face familiarity, working memory and recognition processes (Herzmann et al., 2004; Schweinberger & Burton, 2003; Tanaka et al., 2006). The fact, that the cTBS of the OFA affected the ERPs the strongest in the time-window corresponding to N250, is in line with those previous findings which consider the OFA more than a simple gateway to the face processing network.

First, neuroimaging studies, using uni- and multivariate analytical methods of fMRI data, found convincing evidence of differential activations or activity patterns in the OFA for both experimentally familiarized as well as for famous faces (for a recent review see Kovács, 2020). Second, recent MVPA studies found facial identity specific information in the OFA for familiar faces (Goesaert & Beeck, 2013; Tsantani et al., 2019) and showed that this identity representation is orientation (Anzellotti et al., 2014) and viewpoint independent (Di Visconti Oleggio Castello et al., 2017). Third, the OFA is among the areas which were proposed in the past as a potential cortical source of N250. Although the question is debated currently, previous source localization studies suggested that the N250 may have origins in inferior temporal regions, above all in the fusiform gyrus but including also the inferior occipital gyrus (Kaufmann et al., 2009; Olivares et al., 2018; Schweinberger et al., 2002; Schweinberger et al., 2004), specifically on the right side (Frässle et al., 2016).

Finally, testing the causal role of the rOFA in higher-level face processing we showed that the TMS of the area interferes with the experimental formation of familiarity (Ambrus, Windel et al., 2017) and with familiarity decisions for famous faces in visual (Ambrus, Dotzer et al., 2017), cross-domain priming paradigms (Ambrus et al., 2019) as well as in identity-related semantic associations (Eick et al., 2020). Altogether, these studies suggest that the role of rOFA extends over the traditionally proposed low-level, early, “entry-point” towards higher-level, image-invariant identity processing. Our current study goes further in this line and shows that the electrophysiological signal of differential processing for familiar and unfamiliar faces depends directly on the activations of the rOFA.

### 4.3 Inhibitory cTBS leads to enhanced familiarity effects

cTBS is a physiologically inhibitory TMS protocol, which acts through a process of membrane hyperpolarization and leads to prolonged aftereffects behaviorally (Brückner et al., 2013; Huang et al., 2005; Tarnutzer et al., 2013). We predicted that if the rOFA plays a positive role in the emergence of familiarity representation, then cTBS should reduce the differences of [familiar-unfamilar] ERPs. If, however, these differences are enhanced by cTBS, it likely signals the existence of a more complex background mechanism. We found, somewhat unexpectedly, that cTBS enhanced [familiar-unfamilar] ERP differences, thereby supporting the important role of rOFA in face familiarity processing and at the same time suggesting the existence of an elaborate processing framework for familiar and unfamiliar faces.

Importantly, this is not the first observation of facilitatory effects after inhibitory TMS. cTBS has previously been found to improve both behavioural performance and electrophysiological phenomena in several studies. For example, cTBS improves performance in visual psychophysical tasks (Waterston & Pack, 2010), leads to faster semantic processing of images (Bonnì et al., 2015), enhances the amplitudes of somatosensory evoked potentials (Ishikawa et al., 2007) and results in enhanced brain responses to positive valence words (Roesmann et al., 2019). These studies, together with our current findings, argue against a simple „virtual lesion” account of the effect of cTBS (Brückner et al., 2013) and call for other explanations. In the following, we offer two potential mechanisms for the facilitatory effects of cTBS of the current study.

It has previously been shown that TMS is able to modify the activity of entire networks extensively. For example, Wang et al. (2014), targeted the lateral parietal component of the cortico-hippocampal network with TMS. They found that a multi-session stimulation enhanced the functional connectivity in this network, as well as associative memory performance on the behavioural level, a function dominantly connected to hippocampal processing. Similarly, Handwerker et al. (2020) found that the theta-burst stimulation of the superior temporal sulcus reduced resting-state fMRI connectivity across the entire face processing network. Therefore, cTBS may modulates the inhibitory connections of the rOFA on other cortical sites and this disinhibition leads to the observed facilitations. This hypothesis is in line with the results of a series of TMS experiments of the ventral visual pathway. Pobric et al. (2010) found that repetitive TMS of the occipital pole enhances processing in a semantic association task for words and pictures, a function related to the anterior temporal lobe (ATL), where TMS stimulation delays processing. This explanation assumes that the feed-forward connections of the occipital cortex are inhibitory by nature and that the reduction of this inhibition leads to behavioural improvements (Bonnì et al., 2015). Assuming such a mechanism, the cTBS of the rOFA may enhance the differential processing of familiar faces, by modifying the inhibition of the area on higher-level areas, such as the ATL. Indeed, the role of ATL in familiar face processing has been shown recently by another cTBS study. Chiou and Lambon Ralph (2018) found that cTBS of the right ATL impaired working memory performance selectively for famous faces while leaving unfamiliar face performance intact. Overall, these results suggest a complex functional connectivity pattern across the areas of the face network for the processing of familiar faces and would explain why neuroimaging studies find differential familiarity processings in large portions of this network (for a review see Kovács, 2020).

Another, related, but functionally different, alternative explanation of the facilitatory effects of cTBS is given by Thut and Pascual-Leone (2010). In their review of combined TMS-EEG studies, authors proposed that a larger amplitude ERP component is not necessarily the result of primarily facilitatory effects, but rather it may reflect the onset of secondary, top-down mechanisms, compensating for the inhibition of the TMS. Whether feed-forward disinhibition or the induction of feed-back compensatory excitation is responsible for the facilitatory effects of cTBS will require further, specifically aimed studies.

Another apparently unexpected outcome of the current study is that the stimulation of the right OFA led to a significant cTBS x familiarity interaction over the contralateral occipito-temporal cortex as well. This finding also supports the above explanation that the effect of cTBS modifies the activity of the networks extensively. As the homologue areas of the two hemispheres are always functionally densely connected in the primate brain (Compton, 2002; Frässle et al., 2016), the modulation of one hemisphere may manifests itself over the contralateral homologue as well. The similar modulation of the ERPs over both hemispheres therefore suggests close cooperation of the two OFAs in the processing steps, which are modulated by cTBS.

Independently of the exact neural dynamics of the face processing network under cTBS, our results show that the right occipital face area plays a causal role in the differential processing of familiar and unfamiliar faces and emphasises its role in face recognition processes.

## Acknowledgements

This study was supported by a Deutsche Forschungsgemeinschaft Grant [grant number KO 3918/5-1]. Authors declare no conflict of interest.

